# Microglia States are Susceptible to Senescence and Cholesterol Dysregulation in Alzheimer’s Disease

**DOI:** 10.1101/2024.11.18.624141

**Authors:** Boyang Li, Shaowei Wang, Bilal Kerman, Cristelle Hugo, E Keats Shwab, Chang Shu, Ornit Chiba-Falek, Zoe Arvanitakis, Hussein Yassine

## Abstract

Cellular senescence is a major contributor to aging-related degenerative diseases, including Alzheimer’s disease (AD) but much less is known on the key cell types and pathways driving mechanisms of senescence in the brain. We hypothesized that dysregulated cholesterol metabolism is central to cellular senescence in AD. We analyzed whole transcriptomic data and utilized single-cell RNA seq integration techniques to unveil the convoluted cell-type-specific and sub-cell-type-state-specific senescence pathologies in AD using both ROSMAP and Sea-AD datasets. We identified that microglia are central components to AD associated senescence phenotypes in ROSMAP snRNA-seq data (982,384 nuclei from postmortem prefrontal cortex of 239 AD and 188 non-AD) among non-neuron cell types. We identified that homeostatic, inflammatory, phagocytic, lipid processing and neuronal surveillance microglia states were associated with AD associated senescence in ROSMAP (152,459 microglia nuclei from six regions of brain tissue of 138 early AD, 79 late AD and 226 control subject) and in Sea-AD (82,486 microglia nuclei of 42 dementia, 42 no dementia and 5 reference subjects) via integrative analysis, which preserves the meaningful biological information of microglia cell states across the datasets. We assessed top senescence associated bioprocesses including mitochondrial, apoptosis, oxidative stress, ER stress, endosomes, and lysosomes systems. Specifically, we found that senescent microglia have altered cholesterol related bioprocesses and dysregulated cholesterol. We discovered three gene co-expression modules, which represent the specific cholesterol related senescence transcriptomic signatures in postmortem brains. To validate these findings, the activation of specific cholesterol associated senescence transcriptomic signatures was assessed using integrative analysis of snRNA-seq data from iMGs (microglia induced from iPSCs) exposed to myelin, Abeta, and synaptosomes (56,454 microglia across two replicates of untreated and four treated groups). In vivo cholesterol associated senescence transcriptomic signatures were preserved and altered after treatment with AD pathological substrates in iMGs. This study provides the first evidence that dysregulation of cholesterol metabolism in microglia is a major driver of senescence pathologies in AD. Targeting cholesterol pathways in senescent microglia is an attractive strategy to slow down AD progression.

## Introduction

Alzheimer’s disease (AD) is a progressive neurodegenerative disorder, with aging recognized as the greatest risk factor. Among the hallmarks of aging is cellular senescence, a process characterized by irreversible growth arrest and the gradual decline of cellular function, leading to cell death^1^. Senescent cells experience several impaired biological processes, including DNA damage, abnormalities in endosome-lysosome function, mitochondrial stress, and ER stress^2^. Emerging evidence suggests that senescence in distinct brain cell populations contributes to the neuroinflammatory, and degenerative processes observed in AD. However, the role of microglia, the brain’s resident immune cells, in AD-associated senescence has not been fully elucidated, particularly with regard to the various functional states these cells adopt during disease progression.

AD is associated with the accumulation of senescent glial cells, especially astrocytes, microglia, and oligodendrocytes over time. Glial senescence contributes to neuroinflammation and impaired brain homeostasis, which worsen with aging and AD pathology^3,4^. Microglia, specifically, are highly dynamic and display a range of activation states^5^, such as homeostatic, inflammatory, phagocytic, and lipid processing, each of which can be altered in the context of AD. These functional states enable microglia to play both protective and detrimental roles in the brain, influencing neuroinflammation, synaptic plasticity, and clearance of cellular debris. The susceptibility of these microglial states to senescence in AD, as well as the molecular pathways driving their transition, remains a critical area of investigation. Cholesterol dysregulation is another key feature of AD pathology, with alterations in cholesterol levels and accumulation within cells linked to AD-related changes^6^. Cholesterol homeostasis is crucial for maintaining the integrity of the cell membrane, energy production, and overall cell survival. It is regulated through a balance of processes, such as cholesterol biosynthesis, uptake, efflux, transport within endosome-lysosome compartments, and metabolism^7^. Disruptions in these processes can lead to dangerous cholesterol imbalances, which contribute to cell dysfunction and age-related degeneration^8^.

In this study, we analyzed single-cell RNA sequencing (scRNA-seq) datasets to unravel a comprehensive landscape of cellular senescence across different glial cell types in the brains of patients with AD. Among all cell types, we identified microglia as especially prone to senescence during AD progression. We further investigated cholesterol regulation in relation to AD and cellular senescence, focusing on the different functional states of microglia. We found that both homeostatic and inflammatory microglia were particularly vulnerable to cholesterol-related senescence. Exposing iPSC-derived microglia (iMG) to CNS substrates such as myelin debris or apoptotic cells induced disease associated microglia (DAM) with signatures of cholesterol related senescence. Understanding how senescent microglia contribute to AD pathology may open new avenues for intervention, specifically targeting microglial cholesterol metabolism to ultimately slow disease progression.

## Results

### Heterogeneous expression of cellular senescence markers across glial cell types and AD statuses

The snRNA-seq expression profile of 982,364 nuclei from the prefrontal cortex (PFC) of postmortem brain tissue from 427 participants (Extend Table 1.) in the ROSMAP was re-analyzed, targeting cellular senescence-related genes and pathways^9^. These cells include astrocytes (149,558), microglia (79,188), oligodendrocytes (645,142), oligodendrocyte precursor cells (OPCs; 90,502), and vasculature cells (17,974). Based on the postmortem diagnosis AD, which relies on both neurofibrillary tangles (Braak) and neuritic plaques (CERAD), the study included 239 AD and 188 non-AD participants. To investigate cellular senescence across major glial cell types in the brain, we examined the expression patterns of several canonical cellular senescence markers (pRB/CDKN2A axis: CDKN2A, which respond to DNA damage; TP53/CDKN1A axis: TP53 and CDKN1A, which represses pro-proliferation genes) in astrocytes, microglia, oligodendrocytes, OPCs, and vasculature cells (Fig. 1A). These findings indicate that CDKN1A was widely expressed in all five cell types of interest. In contrast, CDKN2A shows reduced expression levels in cells associated with blood vessels. CDKN2A is more highly expressed in oligodendrocytes and OPCs, whereas TP53 is mainly expressed in astrocytes. To investigate the pathological association between cellular senescence and AD, we compared the percentage of expression (proportion of cells with gene counts ≥ 1 of the respective marker) of canonical cellular senescence markers (pRB/CDKN2A axis: CDKN2A; TP53/CDKN1A axis: ATM, GPNMB, and CDKN1A; DNA damage: H2AFX) between AD and non-AD cells at the subject level (Fig. 1B). These results demonstrate that astrocyte senescence in AD is primarily associated with ATM and H2AFX; microglia: GPNMB; oligodendrocyte: ATM, CDKN2A, and H2AFX; OPC: ATM, and H2AFX; vasculature cell: none. In addition, we annotated the cellular senescence-related genes in the differentially expressed genes between AD and non-AD cells across cell types (Fig. 1C). Consistent with published studies, these annotated upregulated AD-associated genes contributed to senescence by cell type: NFKB1 in astrocytes^10^, SP1 and SERPINE1 in microglia^11^ and NFKB1, ATM, and RELA in oligodendrocytes^12^. As the percentage of expression of cellular senescence markers showed, AD could not be associated with cellular senescence in vasculature cells at the gene level. Taken together, no single gene is sufficient to define senescence across different brain cell types in single-cell RNA sequencing data.

**Fig 1.**
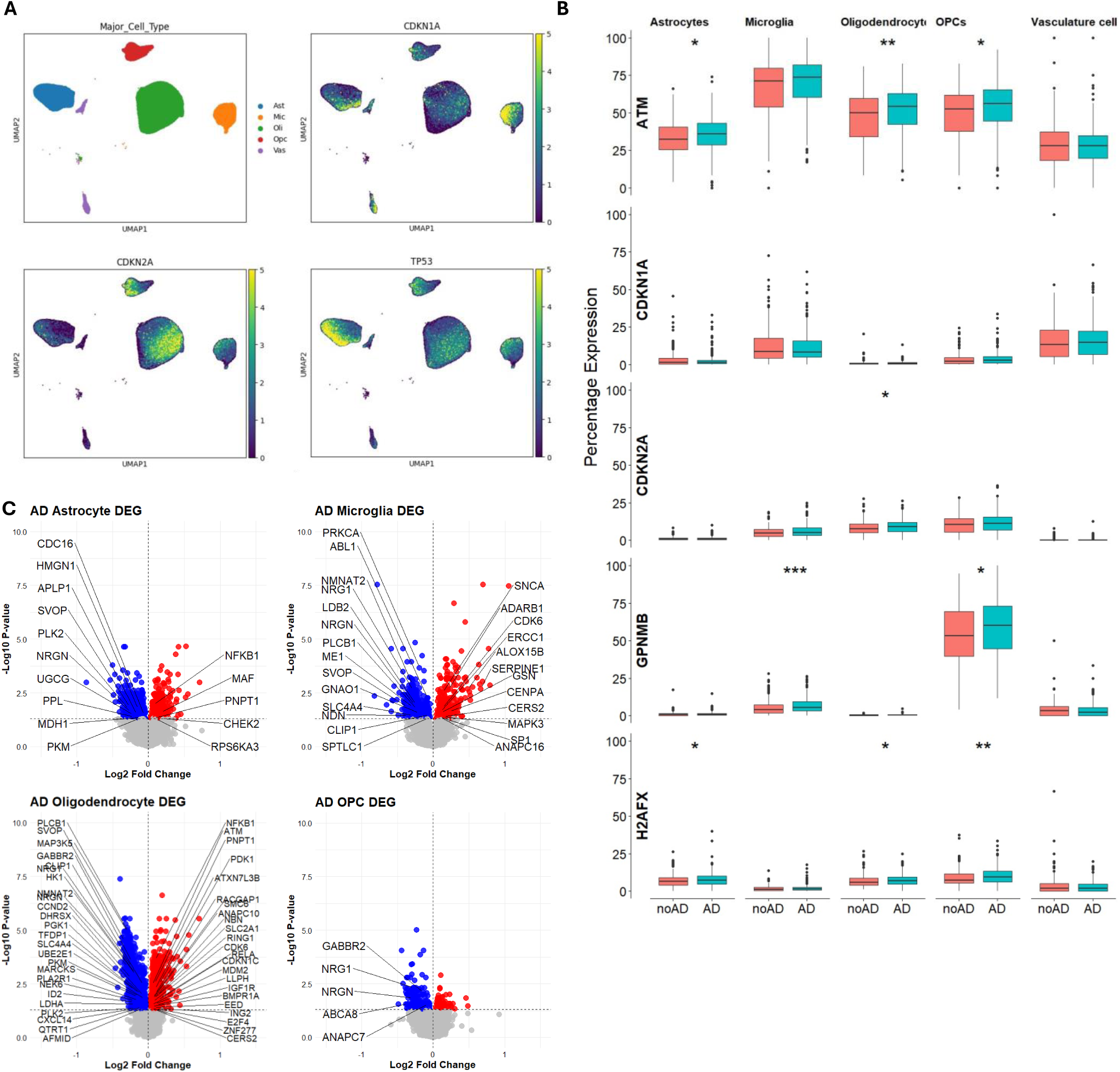
Different cell types have distinct senescence transcriptional signatures. A, Expression pattern of cellular senescence marker genes among non-neuronal cell types (Ast: astrocytes; Mic: microglia; Oli: oligodendrocytes; Opcs: oligodendrocyte progenitor cells; Vas: vasculature cells; Exc and Inh: excitatory and inhibitory neurons, not shown). B, Percentage expression (percentage of nuclei with the gene expression >0) of cellular senescence marker genes among glia cell types. Data shown are median ± quartiles and were analyzed using the Wilcoxon signed-rank test. (*p < 0.05, **p < 0.01, ***p < 0.001, ****p < 0.0001). C, Volcano plots of differential expression genes of AD samples using NEBULA and names of senescence related genes are labelled. Significance is defined as “fdr” adjusted p < 0.05. Redness on the right indicates AD group has a higher expression of the labeled genes.

### AD is Strongly Associated with Cellular Senescence in Microglia

Cellular Senescence is a complex biological process involving multiple features and cell types. To comprehensively understand the association between AD and cellular senescence, we investigated cellular senescence-related pathways, including mitochondrial function, ER stress, oxidative stress, apoptotic signals, and endosome-lysosome trafficking, all of which are altered in brains with AD ^13,14^. We calculated the pathway enrichment scores of these biological processes for each cell using transcriptome-wide profile. We tested whether AD pathology is associated with changes in cellular senescence and related pathways in different non-neuron cell types, preventing pseudoreplication bias from snRNA-seq data. The results demonstrate that microglia and oligodendrocytes are susceptible to cellular senescence associated with AD and its destructive biological process, while vasculature cells, OPCs, and astrocytes are less associated with AD-related cellular senescence (Fig. 2A). In microglia and oligodendrocytes, AD was associated with cellular senescence, mitochondrial abnormalities, and altered apoptotic signals (Fig. 2A). Specifically, the microglial endosome-lysosome system is greatly changed in AD, implying the importance of microglial lysosome-autophagy in exacerbating AD pathologies. Overall, microglia from participants with AD showed a significantly higher pathway score for cellular senescence (Fig. 2B).

**Fig 2.**
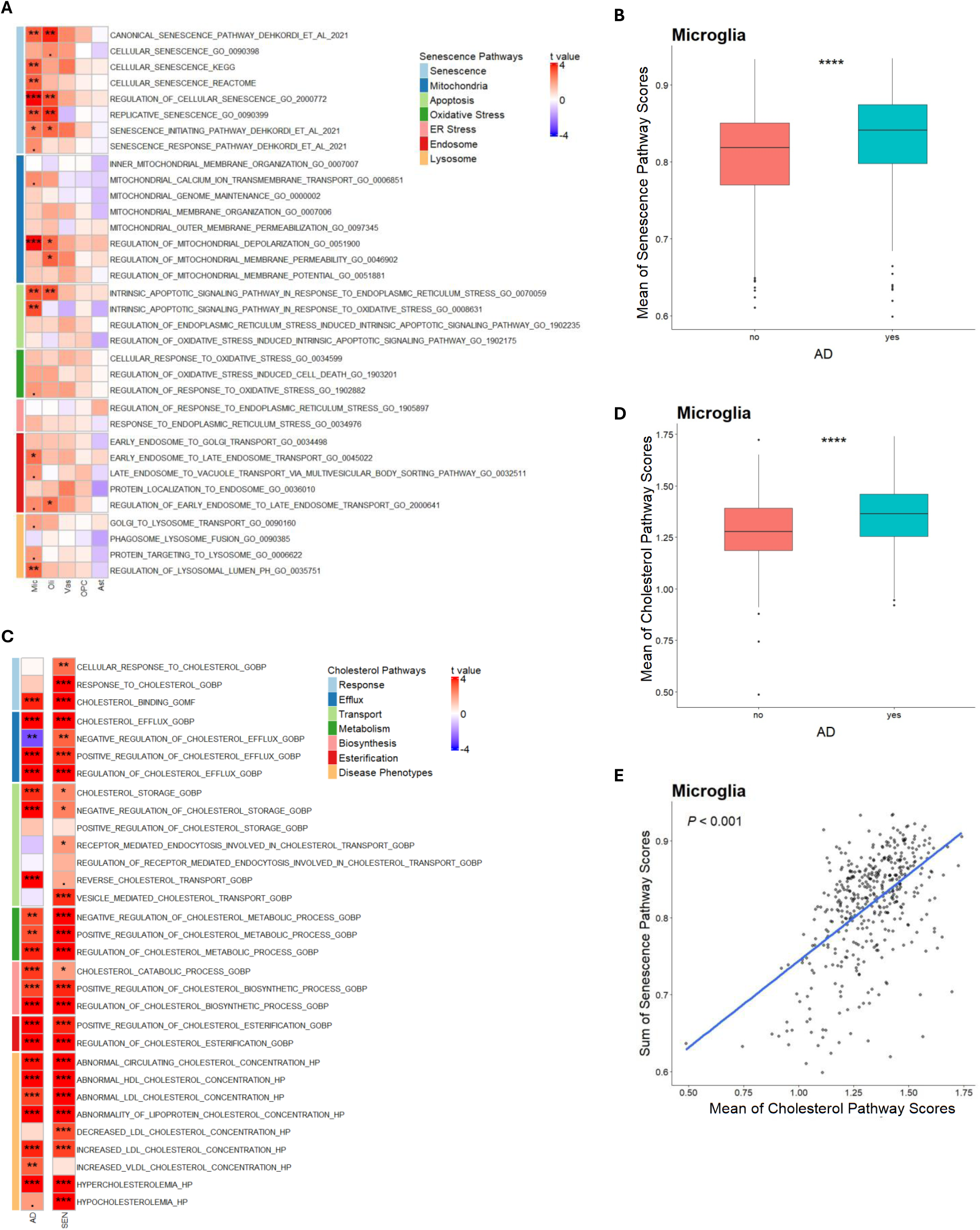
Microglia senescence is important in AD and is associated with cholesterol dysregulation. A, Association of senescence related pathways scores with AD among different cell types using linear mixed effect model. The heatmap represents the t value which is the signed effect size divided by standard derivation. Significance is defined by “fdr” adjusted p value (.p < 0.1, *p < 0.05, **p < 0.01, ***p < 0.001, ****p < 0.0001). B, The individual-level sum of senescence pathway scores are calculated using the average of microglia “Senescence” (the light blue “Senescence” module in the panel 2A) pathways scores for each individual. C, Association of cholesterol related pathways scores with AD or senescent states using linear mixed effect model. The heatmap represents the t value which is the signed effect size divided by standard derivation. D, The individual level of cholesterol pathway scores are calculated using the mean of microglia cholesterol-related (all pathways in the panel 2C) pathways scores together for each individual. Data shown are median ± quartiles and were analyzed using the Wilcoxon signed-rank test. (*p < 0.05, **p < 0.01, ***p < 0.001, ****p < 0.0001). E, The association of senescence and cholesterol related pathway scores, calculated in panel 2B&2D.

Since microglia play important roles in cholesterol homeostasis and myelin phagocytosis in the brain, and these features are impaired in aged or AD subjects^6,15,16^, we hypothesize that senescent microglia have impaired cholesterol regulation machinery, which is associated with worse AD pathology. To test this hypothesis, we investigated whether microglia from AD participants would appear to have similar impairments as senescent microglia in relation to cholesterol-related pathways, including cholesterol response, efflux, storage, transport, metabolism, biosynthesis, esterification, lipid particles, and abnormal cholesterol levels. We found that microglia from AD participants and senescent microglia (senescent state is identified for each cell if its senescence pathway enrichment score is larger than the mean plus 2x standard deviation) have similar alterations across the whole spectrum of cholesterol-related pathways (Fig. 2C). Overall, microglia from AD participants showed a significantly higher pathway score for cholesterol-related features (Fig. 2D). The cholesterol pathway scores were highly correlated with the cellular senescence pathway scores (Fig. 2E). Taken together, microglial senescence is one of the most predominant markers of AD pathology among non-neuronal cell types. Microglial senescence is strongly associated with changes in cholesterol-related bioprocesses and could be a risk factor for AD.

### Homeostatic, Inflammatory, Phagocytic, Lipid Processing and Neuronal Surveillance Microglia are Susceptible to Cholesterol Associated Cellular Senescence in AD in ROSMAP Study

To unveil microglia-state-specific and AD-stage-specific alterations in cellular senescence and cholesterol-related biological processes, we reanalyzed the snRNA-seq expression profile of 164,076 microglia nuclei from six regions of postmortem brain tissue from 424 participants (Extend Table 2.) in ROSMAP. Microglia were classified into 12 clusters (MG0: homeostatic, MG1: neuronal surveillance, MG2: inflammatory I, MG3: ribosome biogenesis, MG4: lipid processing, MG5: phagocytic, MG6: stress-related, MG7: glycolytic, MG8: inflammatory II, MG10: inflammatory III, MG11: antiviral and MG12: cycling) based on their molecular signatures and functions from the original publication^5^. To investigate which microglial cell states are specifically susceptible to cellular senescence, we compared the cellular senescence scores across the 12 identified microglial cell states. The results showed that antiviral microglia had the highest cellular senescence scores, whereas the neuronal surveillance microglia had relatively lower cellular senescence scores (Fig. 3A-3B). To identify which microglial cell states contribute to the development of AD-related cellular senescence, we also tested the cell-state-specific association between AD and cellular senescence during the early and late stages of AD (Fig. 3C-3D). These results demonstrate that homeostatic and inflammatory microglia have cellular senescence scores that change in the early stage of AD, and homeostatic, inflammatory, phagocytic, lipid processing, and neuronal surveillance microglia possess cellular senescence features in the late stage of AD. Overall, these five states of microglia from AD participants showed a significantly higher pathway score for senescence-related features (Fig. 3E). We tested the cell state-specific association between AD and cholesterol homeostasis in the early and late stages of AD. The results demonstrated that inflammatory, lipid processing, and homeostatic microglia have cholesterol-related pathways that change in the early stage of AD, and inflammatory, homeostatic, phagocytic, neuronal surveillance, and lipid processing microglia possess impaired cholesterol homeostasis in the late stage of AD (Fig. 3C-3D). Overall, these five states of microglia from AD participants showed a significantly higher pathway score for cholesterol-related features (Fig. 3F). The cholesterol pathway scores were highly correlated with the cellular senescence pathway scores (Fig. 3G). The results showed that cellular senescence-related and cholesterol-related pathway enrichment scores increased as AD development stages progressed, and cholesterol pathways were correlated with cellular senescence pathways. Taken together, abnormal cholesterol could induce microglial senescence in homeostatic and inflammatory states in AD brains.

**Fig 3.**
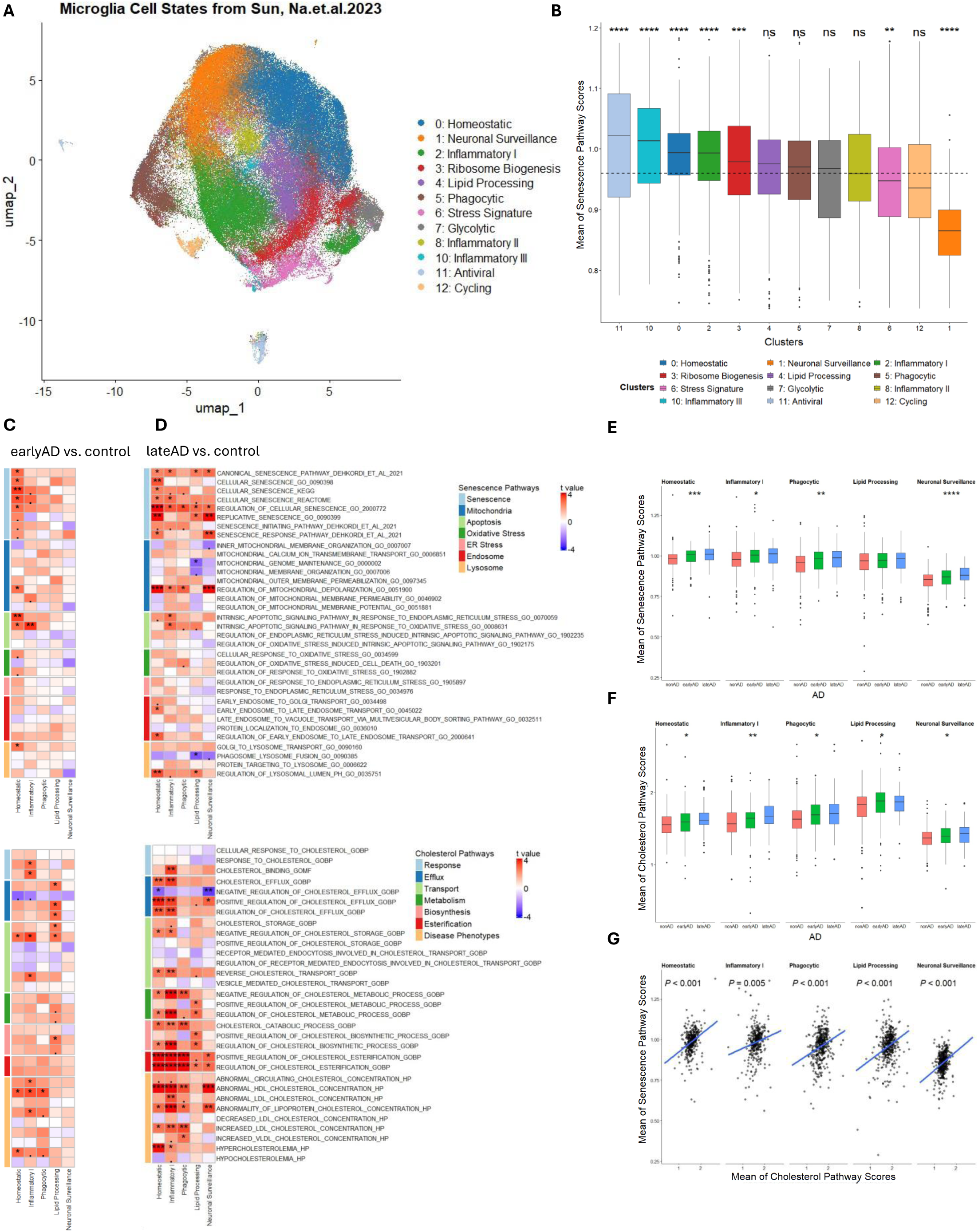
Cell states of microglia are susceptible to cholesterol related senescence in AD. A, UMAP of 164,076 microglia nuclei with annotated microglial states in ROSMAP 6-region snRNA-seq data. B, The individual level of senescence related pathway scores among each microglia cell states calculated using the same method as Fig2. C-D, Association of senescence and cholesterol related pathways scores with AD disease stages (C: earlyAD vs. control; D: lateAD vs. control) in different microglia cell states using linear mixed effect model. The heatmap represents the t value which is the signed effect size divided by standard derivation. Significance is defined by “fdr” adjusted p value (.p < 0.1, *p < 0.05, **p < 0.01, ***p < 0.001, ****p < 0.0001). E-F, The individual level of senescence and cholesterol related pathways scores across AD disease states in homeostatic, inflammatory I, phagocytic, lipid processing and neuronal surveillance microglia calculated using the same method as Fig2. Data shown are median ± quartiles and were analyzed using the Kruskal– Wallis test. (*p < 0.05, **p < 0.01, ***p < 0.001, ****p < 0.0001). G, The association of senescence and cholesterol related pathway scores in homeostatic, inflammatory I, phagocytic, lipid processing and neuronal surveillance microglia.

### Inflammatory Microglia Dominantly Drive Cholesterol Associated Cellular Senescence in Dementia in SEA-AD Study

Single-cell RNA-seq enables full transcriptional characterization of cellular senescence based on microglial cell states. However, it is challenging to unbiasedly compare the association between cellular senescence and impaired cholesterol homeostasis in the development of AD across different datasets because of the variance from different assay methods, technique effects, and biological backgrounds. To improve the robustness of our hypothesis on the association between cellular senescence and impaired cholesterol homeostasis in microglia cell states in AD, we used the “Seurat” embedded “CCA-Integration” function to project microglia from SEA-AD (Extend Table 3, Fig. 4A) onto the cell states from ROSMAP (Fig. 3A), accounting for experimental and biological factors. The results show that inflammatory, homeostatic, and lipid processing microglia are the top three projected microglia in SEA-AD. We tested the cellular senescence-related and cholesterol-related pathway enrichment scores between clinically diagnosed dementia and non-dementia in homeostatic, inflammatory, phagocytic, and lipid-processing microglia (Fig. 4B-4C). The results demonstrate that inflammatory microglia are particularly susceptible to cellular senescence and abnormal cholesterol levels in dementia, while senescence-related and cholesterol-related pathways are not significantly altered if interrogated separately in homeostatic microglia. We tested the summary of cellular senescence-related and cholesterol-related pathway enrichment scores in homeostatic and inflammatory microglia, and determined that cellular senescence-related and cholesterol-related pathway enrichment scores increase with the development of dementia (Fig. 4D-4E), and cholesterol pathways are correlated with cellular senescence pathways (Fig. 4F). Taken together, abnormal cholesterol levels are strongly associated with cellular senescence in inflammatory microglia, whereas this phenotype could be less predominant in homeostatic microglia. This cholesterol-related cellular senescence could contribute to neurodegenerative diseases such as AD.

**Fig 4.**
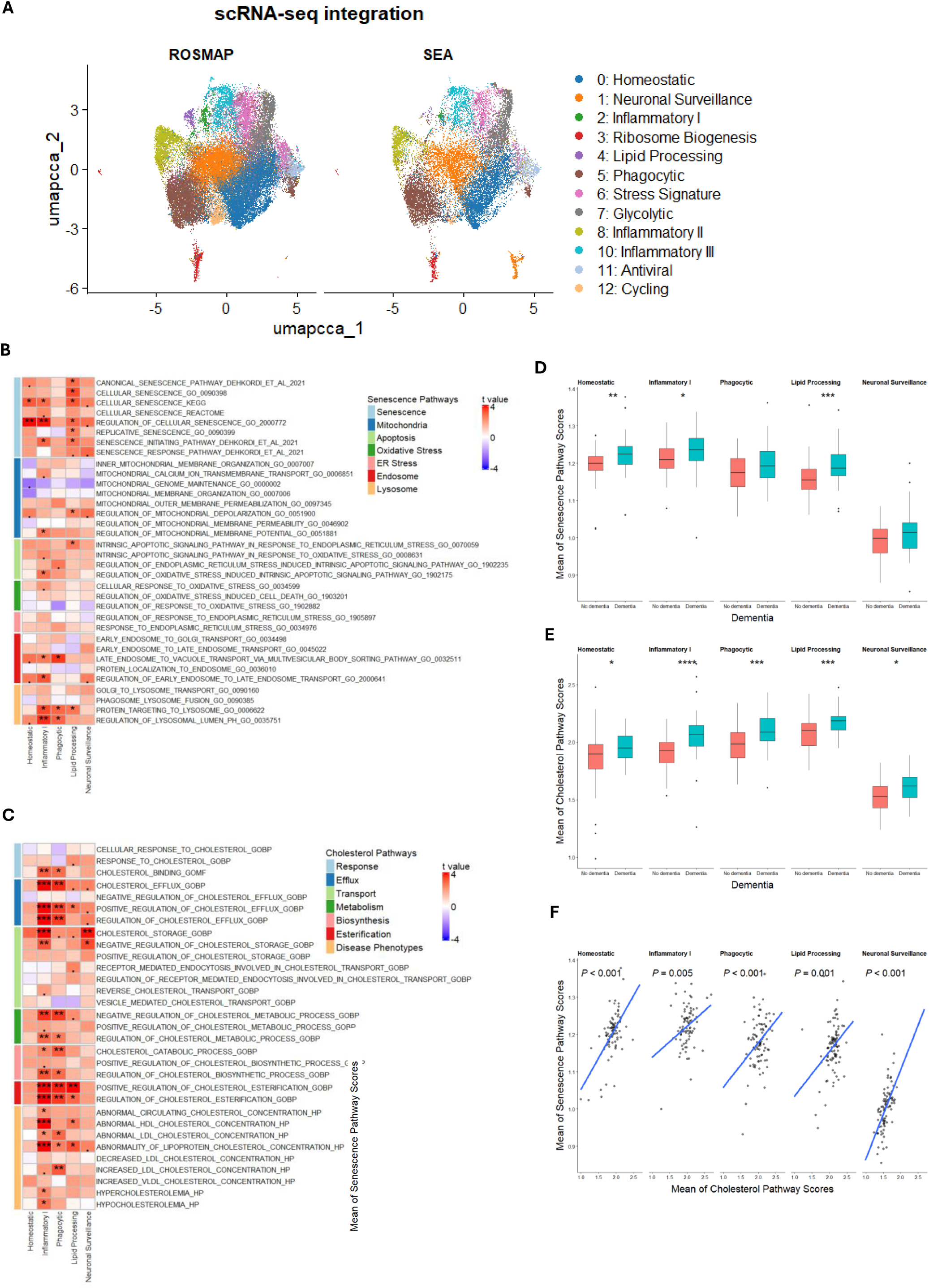
Cell states of microglia from SEA-AD are susceptible to cholesterol related senescence in dementia. A, UMAP of microglia nuclei with annotated microglial states in SEA-AD snRNA-seq data projection from microglia cell state in Fig3. B-C, Association of senescence and cholesterol related pathways scores with dementia in homeostatic, inflammatory I, phagocytic and lipid processing microglia using linear mixed effect model. The heatmap represents the t value which is the signed effect size divided by standard derivation. Significance is defined by “fdr” adjusted p value (.p < 0.1, *p < 0.05, **p < 0.01, ***p < 0.001, ****p < 0.0001). D-E, The individual level of senescence and cholesterol related pathways scores by dementia in homeostatic, inflammatory I, phagocytic and lipid processing microglia calculated using the same method as Fig2. Data shown are median ± quartiles and were analyzed using the Wilcoxon signed-rank test. (*p < 0.05, **p < 0.01, ***p < 0.001, ****p < 0.0001). F, The association of senescence and cholesterol related pathway scores in homeostatic, inflammatory I, phagocytic and lipid processing microglia.

### Cholesterol Associated Senescence Transcriptomic Signatures are Established using ROSMAP snRNA Data

We established co-expression networks of cholesterol-associated senescence genes by creating a list of cholesterol- and senescence-related genes from the Gene Ontology (GO) database and used this list to perform weighted gene co-expression network analysis (WGCNA) to retrieve co-expression modules from the ROSMAP dataset (Fig. 2), representing the specific cholesterol-associated senescence transcriptomic signatures (Fig. 5A). We identified three co-expression modules with altered disease-related characteristics *in vivo* (Fig. 5D-5F). We tested whether these module eigengenes were differentially expressed by overall AD pathology (Fig. 5B) using the “module eigengenes differential expression” function from “Seurat” package, and if these modules are associated with the disease-related variables of interest (Fig. 5C) using the “module-trait association” function. We found that module 1 (blue) and module 3 (brown) eigengenes had higher expression levels in samples with an overall AD pathology. All three modules are associated with probed AD-related variables, indicating that cholesterol-associated senescence signatures are altered in AD.

**Fig 5.**
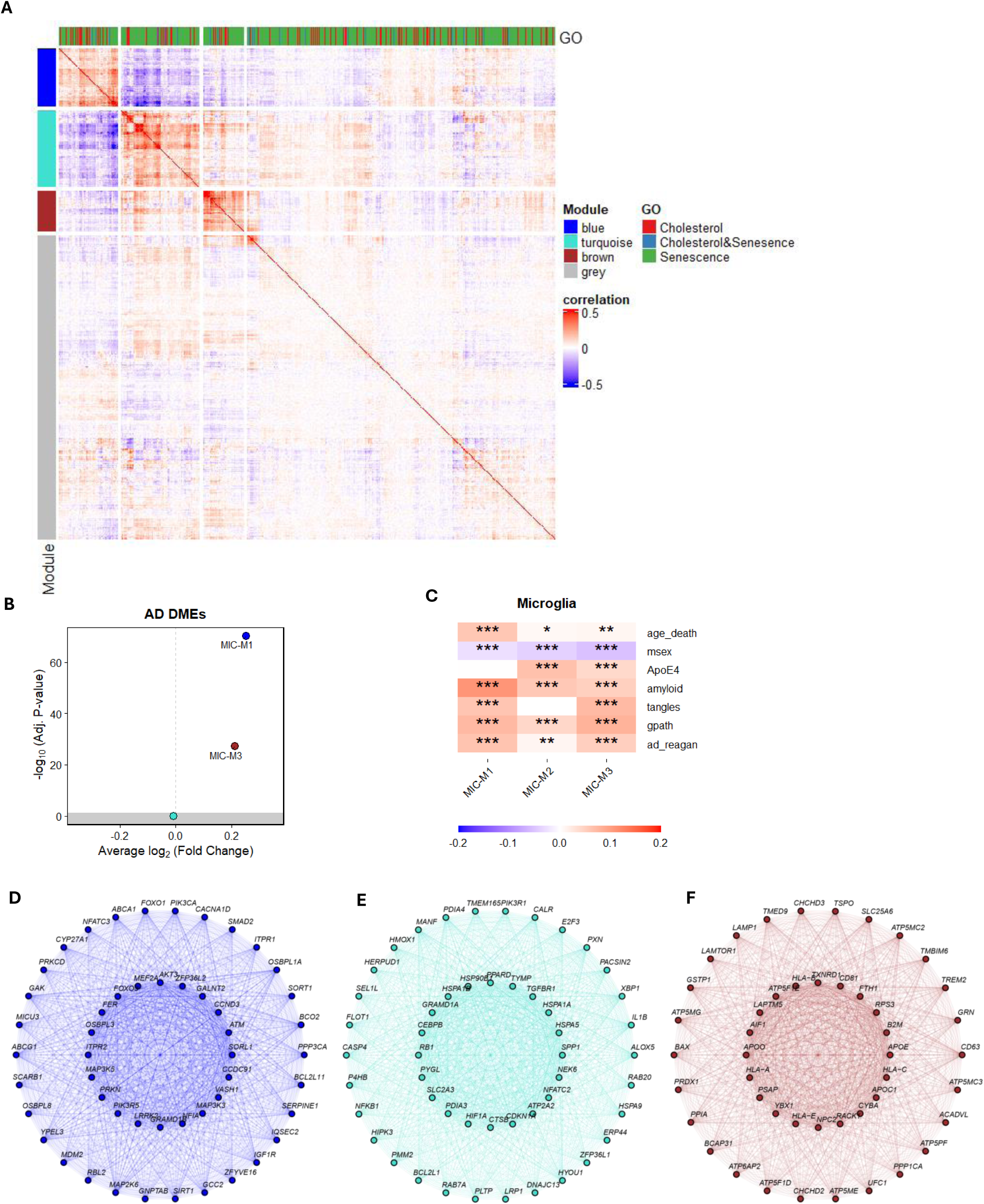
WGCNA gene correlation network analysis shows modules containing senescence and cholesterol related genes in microglia. A, Genes related to senescence and cholesterol are selected for gene correlation network analysis. The heatmap represents the gene-gene correlation and their module identities are labelled blue, turquoise and brown (grey means not correlated). B, Differential expression of module eigengenes by AD. C, Module-traits association plots between the three modules and AD related traits. D-F, Diagrams of the three modules with 20 hub genes and 30 remaining 15 genes.

### Cholesterol Associated Senescence Transcriptomic Signatures are Altered after Treating iMG with CNS Substrates

SnRNA-seq on untreated iPSC-derived microglia (iMGs) or iMGs incubated with synaptosomes (Syn), myelin debris (Myln), apoptotic neurons (Apop) or synthetic amyloid-beta (Ab) fibrils for 24 hours resulted in two clusters of disease associated microglia (DAM, cluster 2 and cluster 8, Fig. 6A-6B) with elevated expression of ABCA1, APOE, GPNMB, LPL (Fig. 6C-6F). We reanalyzed the data of 56,454 single iMG transcriptomes across all conditions for the cholesterol-associated senescence signatures in Fig. 5. The results showed that cholesterol-associated senescence transcriptomic signatures were altered after treatment of iMGs with CNS substrates in DAM clusters 2 and 8 (Fig.6G-6L), indicating that cholesterol-associated senescence phenotypes are important features of AD development.

**Fig 6.**
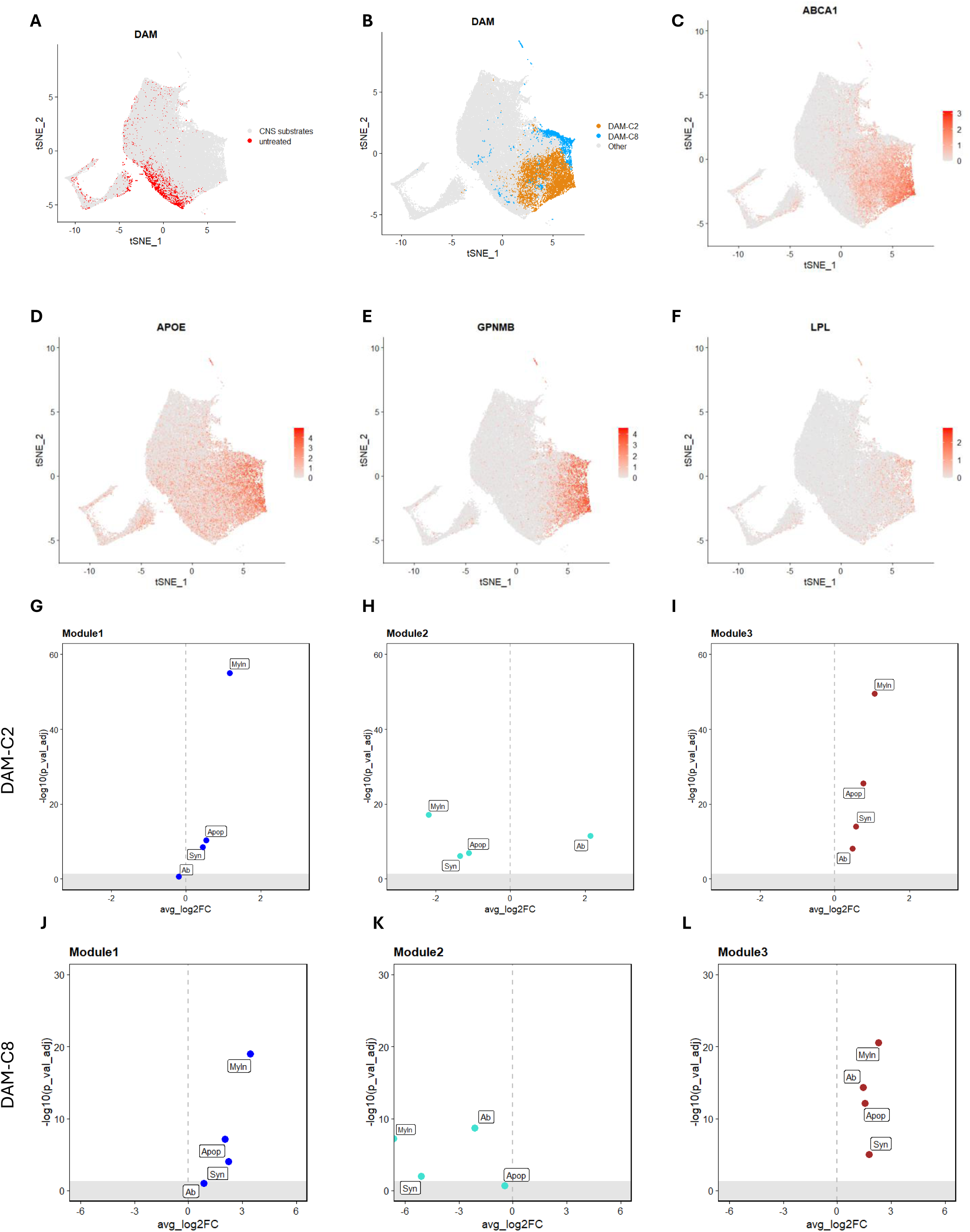
CNS substrate treatments induce DAMs from iMG, displaying signatures of cholesterol related senescence. A-B, DAMs (cluster2 & cluster8) emerge after CNS substrate treatments. C-F, DAM marker expression patterns. G-I, the expression levels of the module eigengenes from Fig5 in DAM cluster2 after CNS substrate treatment. J-L, the expression levels of the module eigengenes from Fig5 in DAM cluster8 after CNS substrate treatment.

## Discussion

In this study, we integrated four independent single-cell transcriptomic datasets to identify AD-associated senescence markers, gene expression differences, biological pathways, and co-expression modules. We annotated the diversity of cell types and cell states within AD-associated senescence molecular signatures at both single-cell and pseudo-bulk levels. Using integrative analysis techniques, we analyzed preserved biological information on senescence signatures and microglial cell states across different datasets (ROSMAP and SEA-AD) and models (human postmortem biospecimens and iPSC-derived microglia). We focused on AD-associated senescence signatures across major non-neuronal cell types in the central nervous system (CNS) and found that microglia from subjects with AD exhibit highly perturbed senescence signatures. This suggests that microglial dysfunction is central to the pathology of AD-related senescence. Further analysis showed that specific microglial states—homeostatic, inflammatory, phagocytic, lipid processing, and neuronal surveillance—are particularly vulnerable to AD-associated senescence. Among these, the inflammatory microglial state dominates the effects of senescence on AD, highlighting microglia as the most susceptible cell type to AD-related senescence, with specific states driving this pathology.

We examined cellular senescence-related pathways including mitochondrial dysfunction, ER stress, oxidative stress, apoptotic signaling, and endosome-lysosome trafficking. Distinct senescence-related processes were observed, including mitochondrial depolarization, stress-induced apoptosis, early to-late endosome transport, and lysosomal pH imbalance. A comparison of molecular perturbations between microglia from AD subjects and senescent microglia revealed that cholesterol-related bioprocesses were highly dysregulated in both groups, suggesting that cholesterol dysregulation is a key driver of AD-associated senescence pathologies. More specifically, our results highlight the importance of microglial cholesterol homeostasis in AD pathology and suggest that targeting cholesterol metabolism in senescent microglia could be a promising therapeutic strategy. The identification of specific microglial states that are more vulnerable to senescence and cholesterol dysregulation provides a clearer understanding of the cell-type and state-specific mechanisms driving AD progression. This opens the door for potential interventions aimed at modulating microglial function and cholesterol homeostasis to mitigate neurodegeneration in AD.

A key finding of our study is the strong correlation between cholesterol dysregulation and microglial senescence in AD. We showed that AD microglia exhibit alterations across multiple cholesterol-related pathways, including cholesterol eplux, storage, metabolism, and biosynthesis. This dysregulation is particularly pronounced in inflammatory and lipid-processing microglial states. These findings were further validated using data from induced microglia (iMGs) treated with CNS substrates, where cholesterol-associated senescence transcriptomic signatures were significantly altered. We identified three distinct gene co-expression network modules that represented cholesterol-associated senescence signatures in AD microglia. Module 1 includes several AD risk genes, such as **ABCA1**, **ABCG1**, and **SORL1**, suggesting that senescent microglia may have altered responses to amyloid-beta. Module 2 contained inflammation-related genes, such as **IL1B**, **NFKB1**, and **CEBPB**, highlighting the importance of inflammatory states in AD-associated senescence. Module 3 contains disease-associated microglial (DAM) genes, such as **APOE**, **B2M**, and **TREM2**, indicating that senescent microglia are key contributors to AD pathology. These modules are also influenced by CNS substrate treatment in iPSC-derived microglia (iMGs), further supporting the role of cholesterol-associated senescence in driving AD development. However, no single gene is sufficient to define senescence across different brain cell types in single-cell RNA sequencing data, possibly because of the limited sequencing depth per cell and the high proportion of zero reads^17^. AD-associated cellular senescence is heterogeneous within cell types and across the brain. Various senescence markers and senescence-related features should be considered to define if a cell is in a senescent state, which will lead to further studies^18^.

In conclusion, our study demonstrates that microglia play a central role in AD-associated cellular senescence, with cholesterol dysregulation acting as a key driver of this process. Future research should focus on developing therapies that target these specific microglial states and their associated cholesterol pathways to slow the progression of AD.

## Methods

### Database

Processed single-nucleus transcriptomic data and ROSMAP metadata were downloaded from the Synapse AD Knowledge Portal (https://www.synapse.org/#!Synapse:syn52293417) with Synapse ID: syn52293433 (Fig. 1&2&5) and syn2580853 (Fig. 3). Source data were collected from 427 (syn52293433) and 443 (syn2580853) subjects from ROSMAP^5,9,19^. Nuclei were isolated from frozen postmortem brain tissues and subjected to droplet-based single-nucleus RNA sequencing (snRNA-seq). Cell types were assigned based on Leiden clustering, marker gene analysis, and comparisons with previously published data in the original publication^5,9^.

Processed single-nucleus transcriptomic data were downloaded from the Seattle Alzheimer’s Disease Brain Cell Atlas (SEA-AD)^20^. Source data were collected from 89 participants (84 subjects and 5 references). Cell types were assigned by hierarchical probabilistic Bayesian mapping to BICCN reference in the original publication^20^. Cell Ranger outputs of iMG (control and CNS substrate treatment) scRNA-seq data were downloaded from Terra. Summary level data are available at https://app.terra.bio/#workspaces/Stevenslab/public_iMGLdatasets.

### Differential Gene Expression (DGE) Analysis Using NEBULA

To perform DGE analysis, we utilized the NEBULA package available in R, which implements a negative binomial mixed-effect model^21^. The model is built to simultaneously account for overdispersion in RNA-seq data and the correlation structure within clusters, ensuring that differentially expressed genes are identified accurately. To annotate senescence related genes, the keyword “SENESCENCE” was used on MsigDB (Molecular Signatures Database) gene sets C2 and C5.

### Gene Set Enrichment Analysis (GSEA, Ucell) for Pathway Scores

The R package GSVA (method option “ssGSEA”) was used to calculate the pathway enrichment score (REACTOME_CELLULAR_SENESCENCE (M27188)) of each cell as the normalized difference in empirical cumulative distribution functions (CDFs) of gene expression ranks^22,23^. The cells were classified as senescent (Fig. 2C: “SEN”) if its senescence pathway enrichment score is larger than mean plus 2x standard deviation. The R package “UCell” was used to calculate the pathway enrichment scores for senescence-related and cholesterol-related pathways^24^. Ucell calculates gene signature scores for snRNA-seq data based on the Mann-Whitney U statistic. Ucell generates pathway enrichment score based on gene rankings for each individual cell. The list of senescence-related pathways was adapted from a study that used single nuclear transcriptomics to characterize senescent cells in AD^2^. The cholesterol-related pathway list was generated from MsigDB using “cholesterol” as the keyword.

### Microglia State Projection

The R package “Seurat” embedded “CCA-Integration” function was used to project the microglia states in ROSMAP onto microglia in SEA-AD^25^. Briefly, ROSMAP microglia were set as the reference dataset with predetermined cell state labels, and SEA-AD microglia were set as the query dataset. Both datasets were log-transformed first, and the top 2000 features were selected before the projection. After several iterations to test the query method for projection, the parameters (reduction = “pcaproject,” reference.reduction = “integrated.cca”, project.query = T) were used to predict microglia states in SEA-AD from ROSMAP microglia states.

### Co-expression network analysis

We calculated the Pearson correlation coefficient between each pair of genes (including senescence-related genes and cholesterol-related genes) across the individuals to generate a correlation matrix (p-value <0.05 in R) using microglia transcriptomic data from Fig. 2. We constructed a co-expression network and extracted the co-expression modules using hdWGCNA from “Seurat.” Module eigengene differential expression (DME), connectivity, and module-trait association were calculated using functions provided by hdWGCNA^26^.

### Statistics

The linear mixed effect model was used with AD pathology as the fixed effect and sample origin as the random effect (parameters and formula in the method), to test if the pathway score was associated with the variables of interest, using the following formula: **pathway score ∼ interested variables + size factor (library size of each cell) + (1|projid (sample id)).** Multiple comparisons were corrected using the false discovery rate (FDR) method. A two-tailed Student’s t-test or a one-way ANOVA followed by a post hoc Tukey’s test with p < 0.05 considered significant were used to test the AD effect.

## Extend Figures

**Extend Table 1.**
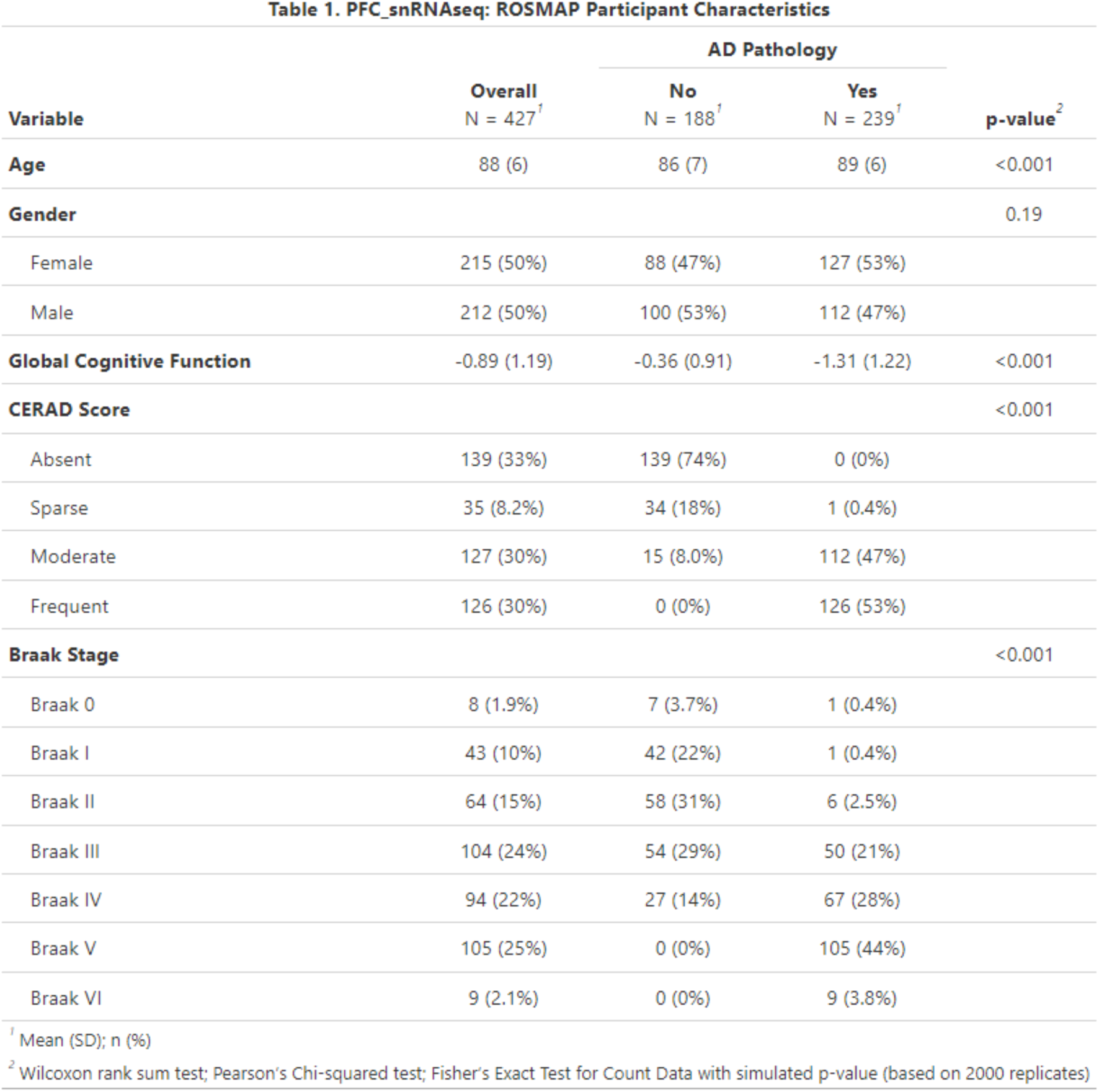
ROSMAP participant characteristics for PFC snRNA-seq data in Fig. 1&2.

**Extend Table 2.** ROSMAP participant characteristics for six-region snRNA-seq data in Fig. 3.

| Variable | Overall<br>N = 424<sup>1</sup> | AD Status |  |  | p-value<sup>2</sup> |
| --- | --- | --- | --- | --- | --- |
|  |  | nonAD<br>N = 219<sup>1</sup> | earlyAD<br>N = 132<sup>1</sup> | lateAD<br>N = 73<sup>1</sup> |  |
| **Age** | 88 (6) | 87 (7) | 89 (6) | 89 (5) | 0.010 |
| **Gender** |  |  |  |  | <0.001 |
| Female | 215 (51%) | 103 (47%) | 60 (45%) | 52 (71%) |  |
| Male | 209 (49%) | 116 (53%) | 72 (55%) | 21 (29%) |  |
| **Global Cognitive Function** | -0.90 (1.19) | -0.34 (0.84) | -1.00 (1.00) | -2.38 (1.09) | <0.001 |
| **CERAD Score** |  |  |  |  | <0.001 |
| Absent | 138 (33%) | 138 (63%) | 0 (0%) | 0 (0%) |  |
| Sparse | 35 (8.3%) | 30 (14%) | 5 (3.8%) | 0 (0%) |  |
| Moderate | 126 (30%) | 42 (19%) | 71 (54%) | 13 (18%) |  |
| Frequent | 125 (29%) | 9 (4.1%) | 56 (42%) | 60 (82%) |  |
| **Braak Stage** |  |  |  |  | <0.001 |
| Braak 0 | 8 (1.9%) | 7 (3.2%) | 1 (0.8%) | 0 (0%) |  |
| Braak I | 43 (10%) | 41 (19%) | 2 (1.5%) | 0 (0%) |  |
| Braak II | 63 (15%) | 52 (24%) | 11 (8.3%) | 0 (0%) |  |
| Braak III | 103 (24%) | 78 (36%) | 25 (19%) | 0 (0%) |  |
| Braak IV | 94 (22%) | 41 (19%) | 52 (39%) | 1 (1.4%) |  |
| Braak V | 104 (25%) | 0 (0%) | 41 (31%) | 63 (86%) |  |
| Braak VI | 9 (2.1%) | 0 (0%) | 0 (0%) | 9 (12%) |  |

<sup>1</sup> Mean (SD); n (%)
<sup>2</sup> Kruskal-Wallis rank sum test; Pearson's Chi-squared test; Fisher's Exact Test for Count Data with simulated p-value (based on 2000 replicates)

**Extend Table 3.** SEA-AD participant characteristics for snRNA-seq data in Fig. 4.

| Variable | Overall<br>N = 84<sup>1</sup> | Cognitive Status |  | p-value<sup>2</sup> |
| --- | --- | --- | --- | --- |
|  |  | No dementia<br>N = 42<sup>1</sup> | Dementia<br>N = 42<sup>1</sup> |  |
| **Age** | 89 (8) | 90 (8) | 88 (8) | 0.37 |
| **Gender** |  |  |  | 0.50 |
| Female | 51 (61%) | 24 (57%) | 27 (64%) |  |
| Male | 33 (39%) | 18 (43%) | 15 (36%) |  |
| **CERAD Score** |  |  |  | <0.001 |
| Absent | 16 (19%) | 14 (33%) | 2 (4.8%) |  |
| Frequent | 30 (36%) | 6 (14%) | 24 (57%) |  |
| Moderate | 26 (31%) | 12 (29%) | 14 (33%) |  |
| Sparse | 12 (14%) | 10 (24%) | 2 (4.8%) |  |
| **Braak Stage** |  |  |  | <0.001 |
| Braak 0 | 2 (2.4%) | 2 (4.8%) | 0 (0%) |  |
| Braak II | 4 (4.8%) | 2 (4.8%) | 2 (4.8%) |  |
| Braak III | 6 (7.1%) | 4 (9.5%) | 2 (4.8%) |  |
| Braak IV | 23 (27%) | 19 (45%) | 4 (9.5%) |  |
| Braak V | 34 (40%) | 14 (33%) | 20 (48%) |  |
| Braak VI | 15 (18%) | 1 (2.4%) | 14 (33%) |  |

<sup>1</sup> Mean (SD); n (%)
<sup>2</sup> Wilcoxon rank sum test; Pearson's Chi-squared test; Fisher's Exact Test for Count Data with simulated p-value (based on 2000 replicates)

## Notes

**Funding** This work was funded in part by the National Institutes of Health/National Institute on Aging (NIH/NIA), RF1AG076124, R01AG055770, R01AG067063, R01AG054434, R21AG056518, and P30AG066530 to H.N.Y.; R01AG082362 and P30AG10161, P30AG72975, and R01AG15819 to R.O.S. and R01 AG057522 and RF1 AG077695 to OC-F]), the Alzheimer’s Drug Discovery Foundation (ADDF) (GC-201711–2014197 to H.N.Y.), and donations from the Vranos and Tiny Foundations and Ms. Lynne Nauss to H.N.Y

### Competing Interest Statement

The authors have declared no competing interest.

## References

1. Dodig, S., Čepelak, I., and Pavić, I. (2019). Hallmarks of senescence and aging. Biochem Med (Zagreb) 29, 030501. 10.11613/BM.2019.030501.

2. Fancy, N.N., Smith, A.M., Caramello, A., Tsartsalis, S., Davey, K., Muirhead, R.C.J., McGarry, A., Jenkyns, M.H., Schneegans, E., Chau, V., et al. (2024). Characterisation of premature cell senescence in Alzheimer’s disease using single nuclear transcriptomics. Acta Neuropathol 147, 78. 10.1007/s00401-024-02727-9.

3. Silva, M.V.F., Loures, C.M.G., Alves, L.C.V., de Souza, L.C., Borges, K.B.G., and Carvalho, M.D.G. (2019). Alzheimer’s disease: risk factors and potentially protective measures. J Biomed Sci 26, 33. 10.1186/s12929-019-0524-y.

4. Lau, V., Ramer, L., and Tremblay, M. (2023). An aging, pathology burden, and glial senescence build-up hypothesis for late onset Alzheimer’s disease. Nat Commun 14, 1670. 10.1038/s41467-023-37304-3.

5. Sun, N., Victor, M.B., Park, Y.P., Xiong, X., Scannail, A.N., Leary, N., Prosper, S., Viswanathan, S., Luna, X., Boix, C.A., et al. (2023). Human microglial state dynamics in Alzheimer’s disease progression. Cell 186, 4386–4403.e4329. 10.1016/j.cell.2023.08.037.

6. Haney, M.S., Pálovics, R., Munson, C.N., Long, C., Johansson, P.K., Yip, O., Dong, W., Rawat, E., West, E., Schlachetzki, J.C.M., et al. (2024). APOE4/4 is linked to damaging lipid droplets in Alzheimer’s disease microglia. Nature 628, 154–161. 10.1038/s41586-024-07185-7.

7. Zhang, J., and Liu, Q. (2015). Cholesterol metabolism and homeostasis in the brain. Protein Cell 6, 254–264. 10.1007/s13238-014-0131-3.

8. Wang, S., Li, B., Cai, Z., Hugo, C., Li, J., Sun, Y., Qian, L., Remaley, A.T., Tcw, J., Chui, H.C., et al. (2024). Cellular senescence induced by cholesterol accumulation is mediated by lysosomal ABCA1 in APOE4 and AD. Res Sq. 10.21203/rs.3.rs-4373201/v1.

9. Mathys, H., Peng, Z., Boix, C.A., Victor, M.B., Leary, N., Babu, S., Abdelhady, G., Jiang, X., Ng, A.P., Ghafari, K., et al. (2023). Single-cell atlas reveals correlates of high cognitive function, dementia, and resilience to Alzheimer’s disease pathology. Cell 186, 4365–4385.e4327. 10.1016/j.cell.2023.08.039.

10. Jong Huat, T., Camats-Perna, J., Newcombe, E.A., Onraet, T., Campbell, D., Sucic, J.T., Martini, A., Forner, S., Mirzaei, M., Poon, W., et al. (2024). The impact of astrocytic NF-κB on healthy and Alzheimer’s disease brains. Sci Rep 14, 14305. 10.1038/s41598-024-65248-1.

11. Gross, P.S., Laforet, V.D., Manavi, Z., Zia, S., Lee, S.H., Shults, N., Selva, S., Alvarez, E., Plemel, J.R., Schafer, D.P., and Huang, J.K. (2024). Senescent-like microglia limit remyelination through the senescence associated secretory phenotype. bioRxiv. 10.1101/2024.05.23.595605.

12. Schlett, J.S., Mettang, M., Skaf, A., Schweizer, P., Errerd, A., Mulugeta, E.A., Hein, T.M., Tsesmelis, K., Tsesmelis, M., Büttner, U.F.G., et al. (2023). NF-κB is a critical mediator of post-mitotic senescence in oligodendrocytes and subsequent white matter loss. Mol Neurodegener 18, 24. 10.1186/s13024-023-00616-5.

13. Jagtap, Y.A., Kumar, P., Kinger, S., Dubey, A.R., Choudhary, A., Gutti, R.K., Singh, S., Jha, H.C., Poluri, K.M., and Mishra, A. (2023). Disturb mitochondrial associated proteostasis: Neurodegeneration and imperfect ageing. Front Cell Dev Biol 11, 1146564. 10.3389/fcell.2023.1146564.

14. Butterfield, D.A., and Halliwell, B. (2019). Oxidative stress, dysfunctional glucose metabolism and Alzheimer disease. Nat Rev Neurosci 20, 148–160. 10.1038/s41583-019-0132-6.

15. McNamara, N.B., Munro, D.A.D., Bestard-Cuche, N., Uyeda, A., Bogie, J.F.J., Hoffmann, A., Holloway, R.K., Molina-Gonzalez, I., Askew, K.E., Mitchell, S., et al. (2023). Microglia regulate central nervous system myelin growth and integrity. Nature 613, 120–129. 10.1038/s41586-022-05534-y.

16. Byrns, C.N., Perlegos, A.E., Miller, K.N., Jin, Z., Carranza, F.R., Manchandra, P., Beveridge, C.H., Randolph, C.E., Chaluvadi, V.S., Zhang, S.L., et al. (2024). Senescent glia link mitochondrial dysfunction and lipid accumulation. Nature 630, 475–483. 10.1038/s41586-024-07516-8.

17. Zyla, J., Papiez, A., Zhao, J., Qu, R., Li, X., Kluger, Y., Polanska, J., Hatzis, C., Pusztai, L., and Marczyk, M. (2023). Evaluation of zero counts to better understand the discrepancies between bulk and single-cell RNA-Seq platforms. Comput Struct Biotechnol J 21, 4663–4674. 10.1016/j.csbj.2023.09.035.

18. Cohn, R.L., Gasek, N.S., Kuchel, G.A., and Xu, M. (2023). The heterogeneity of cellular senescence: insights at the single-cell level. Trends Cell Biol 33, 9–17. 10.1016/j.tcb.2022.04.011.

19. De Jager, P.L., Ma, Y., McCabe, C., Xu, J., Vardarajan, B.N., Felsky, D., Klein, H.U., White, C.C., Peters, M.A., Lodgson, B., et al. (2018). A multi-omic atlas of the human frontal cortex for aging and Alzheimer’s disease research. Sci Data 5, 180142. 10.1038/sdata.2018.142.

20. Gabitto, M.I., Travaglini, K.J., Rachleff, V.M., Kaplan, E.S., Long, B., Ariza, J., Ding, Y., Mahoney, J.T., Dee, N., Goldy, J., et al. (2024). Integrated multimodal cell atlas of Alzheimer’s disease. Nat Neurosci. 10.1038/s41593-024-01774-5.

21. He, L., Davila-Velderrain, J., Sumida, T.S., Hafler, D.A., Kellis, M., and Kulminski, A.M. (2021). NEBULA is a fast negative binomial mixed model for differential or co-expression analysis of large-scale multi-subject single-cell data. Commun Biol 4, 629. 10.1038/s42003-021-02146-6.

22. Hänzelmann, S., Castelo, R., and Guinney, J. (2013). GSVA: gene set variation analysis for microarray and RNA-seq data. BMC Bioinformatics 14, 7. 10.1186/1471-2105-14-7.

23. Barbie, D.A., Tamayo, P., Boehm, J.S., Kim, S.Y., Moody, S.E., Dunn, I.F., Schinzel, A.C., Sandy, P., Meylan, E., Scholl, C., et al. (2009). Systematic RNA interference reveals that oncogenic KRAS-driven cancers require TBK1. Nature 462, 108–112. 10.1038/nature08460.

24. Andreatta, M., and Carmona, S.J. (2021). UCell: Robust and scalable single-cell gene signature scoring. Comput Struct Biotechnol J 19, 3796–3798. 10.1016/j.csbj.2021.06.043.

25. Stuart, T., Butler, A., Hoffman, P., Hafemeister, C., Papalexi, E., Mauck, W.M., Hao, Y., Stoeckius, M., Smibert, P., and Satija, R. (2019). Comprehensive Integration of Single-Cell Data. Cell 177, 1888–1902.e1821. 10.1016/j.cell.2019.05.031.

26. Morabito, S., Reese, F., Rahimzadeh, N., Miyoshi, E., and Swarup, V. (2023). hdWGCNA identifies co-expression networks in high-dimensional transcriptomics data. Cell Rep Methods 3, 100498. 10.1016/j.crmeth.2023.100498.

